# Mitotic adaptations shape acquired resistance and vulnerabilities to KIF18A inhibition in cancer

**DOI:** 10.64898/2026.08.07.743460

**Authors:** Sjoerd J. Klaasen, Kruno Vukušić, Michael R. van Gerven, Haico van Attikum, Iva M. Tolić, Martijn S. Luijsterburg

**Affiliations:** Department of Human Genetics, Leiden University Medical Center, Leiden, The Netherlands; Division of Molecular Biology, Ruđer Bošković Institute, Zagreb, Croatia

## Abstract

KIF18A inhibition selectively kills cancer cells by inducing chromosome alignment defects that activate a spindle assembly checkpoint (SAC)-dependent mitotic arrest. The mechanisms by which initially sensitive cancer cells acquire resistance to KIF18A inhibitors (KIF18Ai), the vulnerabilities of resistant cells, and the initial genetic determinants of KIF18Ai sensitivity remain poorly understood. Using orthogonal CRISPR–Cas9 screening and long-term drug adaptation approaches, we identify two convergent resistance mechanisms. Resistant cells either partially override the SAC, permitting mitotic exit despite chromosome misalignment, or adapt spindle microtubule dynamics to restore chromosome alignment in the absence of KIF18A. Both strategies sustain mitotic progression without inducing KIF18Ai dependence or increasing sensitivity to other mitotic perturbations. In contrast, reduced activity of the mitotic exit regulators PP2A or APC/C enhances KIF18Ai sensitivity. Accordingly, Mps1 inhibitor-driven APC/C mutations enhance responsiveness to KIF18A inhibition. In conclusion, resistance to KIF18A inhibition emerges rapidly, yet distinct genetic contexts create exploitable vulnerabilities to KIF18Ai.

## Introduction

Accurate chromosome segregation during mitosis is essential for preserving genomic stability. Kinesin family member 18A (KIF18A), a kinesin-8 motor protein, plays a central role in this process by limiting microtubule growth at kinetochores, thereby promoting proper chromosome alignment^1, 2^. When KIF18A activity is lost, mitotic spindles become severely distorted and chromosomes fail to align, triggering activation of the spindle assembly checkpoint (SAC)^3, 4^. This leads to a prolonged and ultimately toxic mitotic arrest. Recent work has identified KIF18A as an attractive therapeutic target for selectively eliminating highly aneuploid or whole-genome doubled (WGD) cancer cells^5–7^. These cells are thought to rely more heavily on intact KIF18A function, as increased chromosome numbers raise the likelihood of chromosome misalignment and sustained SAC activation. In addition, other sensitizing contexts linked to KIF18A deficiency have been described, including reduced basal activity of the anaphase-promoting complex/cyclosome (APC/C)^8^. Together, these observations suggest that sensitivity to loss of KIF18A activity is driven by multiple underlying mechanisms.

KIF18A-specific inhibitors suitable for clinical use have only become available in recent years^8–10^ and are currently being evaluated in phase 1 and 2 clinical trials across a broad range of cancer types (Identifier NCT04293094, NCT05902988, NCT06084416 and NCT06799065). However, as with many anti-mitotic therapies, the emergence of acquired resistance is likely to pose a major challenge to long-term treatment efficacy^11^. Although direct evidence for resistance to KIF18A inhibition is lacking, several observations point to plausible resistance mechanisms. Firstly, experimentally introduced point mutations in the KIF18A gene can prevent inhibitor binding without impairing KIF18A function^8^. Secondly, because KIF18A loss preferentially kills cells with high chromosome numbers^5, 7^, reducing chromosome content could restore normal cell division. Thirdly, weakening the SAC improves viability in KIF18A-deficient cells^4, 8^. Finally, mitotic defects caused by KIF18A loss can be alleviated by restoring normal microtubule dynamics, either through disruption of other spindle regulators such as HSET^8^, HAUS8 or NuMA^12^, or by stabilizing microtubules with low-dose taxol^12^

Responses to KIF18A inhibitors (KIF18Ai) vary widely both between and within tumor types^8, 10, 13^. While some cancer cells undergo prolonged mitotic arrest followed by cell death upon KIF18A inhibition, others escape with minimal consequences, pointing to the existence of intrinsic sensitivity and resistance mechanisms. The basis of this variability remains poorly understood. Clarifying both intrinsic and acquired determinants of KIF18Ai response will be essential for developing rational therapeutic strategies and improving patient stratification. In this study, we investigate the mechanisms underlying resistance to KIF18A inhibition and identify vulnerabilities in KIF18Ai-naïve cancers that may enable more effective treatment approaches.

## Results

### Cancer cells rapidly acquire resistance to KIF18A inhibition

To test whether cancer cells can develop resistance to KIF18A inhibitors, we treated seven cancer cell lines (HT-29; colorectal adenocarcinoma, Caco-2; colorectal adenocarcinoma, U2OS; osteosarcoma, Hela; cervical cancer, MDA-MB-231; breast carcinoma, H1299; lung adenocarcinoma and HCT116; colorectal adenocarcinoma,) with 250 nM of the highly selective KIF18A inhibitor sovilnesib (Figure 1a) (NCT06084416). At this concentration, KIF18A was efficiently inhibited, as indicated by its relocalization from kinetochores to spindle poles in all tested cell lines (Figure S1a-c), which is in agreement with a previous^14^⍰. Over the course of several weeks, a subset of these cell lines showed a pronounced reduction in proliferation together with a clear increase in the fraction of mitotic cells. Notably, these effects became less severe with continued culture, suggesting that the cells had adapted and developed resistance. To confirm this, we quantified mitotic timing using brightfield live-cell microscopy after at least six weeks of continuous exposure to either DMSO or KIF18Ai, which we refer to as naïve and exposed cells, respectively. Naïve HT-29, Caco-2 and U2OS cells displayed a severe to intermediate mitotic arrest immediately upon KIF18Ai treatment (Figure 1b–c), whereas this arrest was strongly attenuated in exposed cells. In contrast, naïve Hela, MDA-MB-231, H1299 and HCT116 cells exhibited a much milder mitotic arrest. Even in these lines, exposed cells spent slightly less time in mitosis than their naïve counterparts, indicating that mild resistance also emerged over time. Consistent with these observations, cell viability measurements using CellTiter-Glo confirmed the development of resistance to KIF18Ai in some of the exposed cell lines (Figure 1d). Together, these results demonstrate that cancer cell lines can acquire resistance to KIF18A inhibition during prolonged treatment.

**Figure 1.**
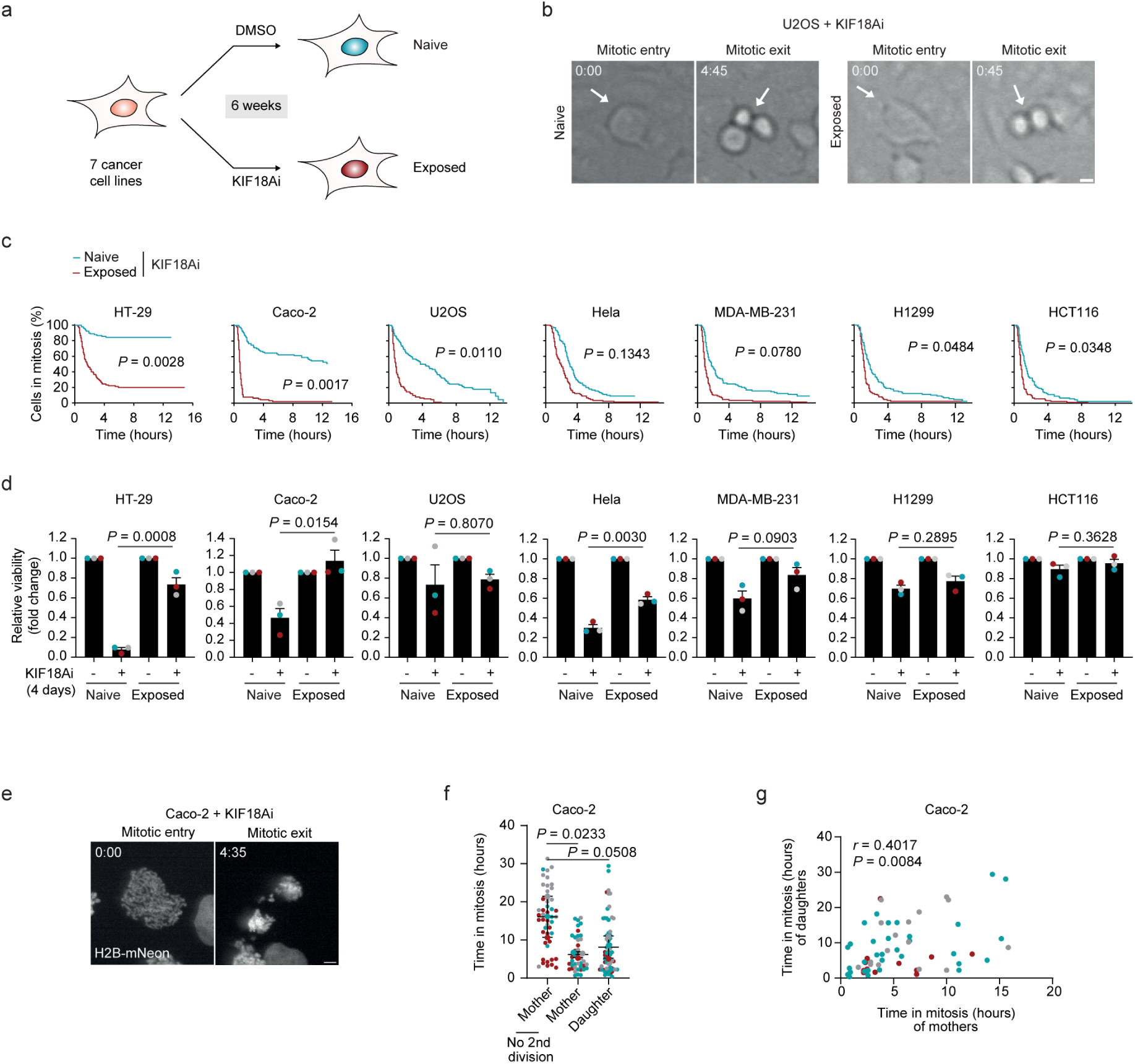
Cancer cells acquire resistance to KIF18Ai over time. (a) Schematic of the generation of seven naïve and KIF18Ai-exposed cancer cell-lines. (b) Representative live-cell imaging stills, and (c) quantification of time in mitosis of naïve and KIF18Ai-exposed cancer cells treated with KIF18Ai. Experiment was performed in triplicate (two-way unpaired t-test; *n* = 90 cells). Scale bar = 5 µm. (d) Cell viability using CellTiter-Glo of naïve and exposed cancer cells after 4 days of DMSO or KIF18Ai treatment. Experiment was performed in triplicate (mean ± s.d., two-tailed unpaired t-test). (e) Example stills and (f, g) quantification of the time in mitosis of naïve Caco-2 cells treated with KIF18Ai. Experiment was performed in triplicate. (f) Mean ± s.d., one-way ANOVA with Dunnett’s multiple comparisons test; *n* = 48, 42 and 67, respectively. (g) Two-tailed Pearson correlation coefficient. Scale bar = 5 µm.

We next wondered when resistance to KIF18Ai arises, and whether a pre-existing subset of cells is intrinsically less sensitive. Although prolonged mitotic arrest was common in the most sensitive lines, a small subset of cells consistently arrested for only a few hours (Figure 1c). This raised the possibility that pre-existing genetic or epigenetic resistance mechanisms might be present. To explore this further, we performed long-term live-cell imaging of naïve Caco-2 H2B-mNeon cells treated with KIF18Ai and followed individual cells across multiple divisions. Cells that proceeded into a second mitosis experienced a significantly shorter first mitosis (6.18 ± 0.78 hours) than cells that did not divide again (16.10 ± 5.39 hours) (Figure 1e–f). In addition, the average duration of second mitoses (8.16 ± 1.92 hours) was shorter than the average duration of all first mitoses (11.43 ± 3.92 hours) and correlated with the length of the first mitosis (Figure 1g). These findings indicate that a pre-existing subpopulation of mildly KIF18Ai-sensitive cells is rapidly selected early during exposure to KIF18Ai.

We then sought to identify the mechanisms underlying resistance to KIF18Ai. One possibility is that KIF18A is no longer effectively inhibited, for example due to mutations in the *KIF18A* gene, increased KIF18A expression, or enhanced drug efflux of the inhibitor. To test this, we examined KIF18A localization on the mitotic spindle. If inhibition was impaired in adapted cells, KIF18A would be expected to remain near kinetochores. As expected, KIF18A localized near kinetochores in both naïve and exposed cells in the absence of inhibitor (Figure S2a). Upon KIF18Ai treatment, however, KIF18A relocalized toward spindle poles in both naïve and exposed cells, indicating that the inhibitor remained effective. This conclusion was further supported by Sanger sequencing of the *KIF18A* gene (Figure S2b), western blot analysis of KIF18A protein levels (Figure S2c), and treatment with a ten-fold higher concentration of the inhibitor (Figure S2d). Another potential explanation is that cancer cells reduce their chromosome number to escape KIF18Ai toxicity, since highly aneuploid and whole-genome doubled cells are particularly sensitive to reduced KIF18A activity^5, 7^. However, flow cytometry-based cell cycle analysis revealed that naïve and exposed cells had comparable DNA content (Figure S2e), ruling out changes in ploidy as a resistance mechanism. Taken together, we conclude that KIF18Ai resistance does not arise from impaired inhibitor binding to KIF18A or by alterations in ploidy.

### A CRISPR-Cas9 screen identifies pathways that modulate KIF18Ai sensitivity

To discover additional mechanisms that influence sensitivity to KIF18A inhibition, we performed a CRISPR-Cas9 knockout screen in RPE1-hTERT TP53 knockout cells treated with either DMSO or KIF18Ai. Although RPE1-hTERT cells are non-cancerous and near-diploid, they retain moderate sensitivity to KIF18Ai (Figure S3a), making them well suited for identifying both genetic sensitizers and desensitizers. This genetic screen uncovered 388 genes whose loss significantly increased or decreased KIF18Ai sensitivity (Figure 2a), representing nearly a ten-fold increase in hits compared with a previous report^8^. Over-representation analysis showed that many of these genes are involved in mitosis-related biological processes (Figure 2b), supporting their functional relevance.

**Figure 2.**
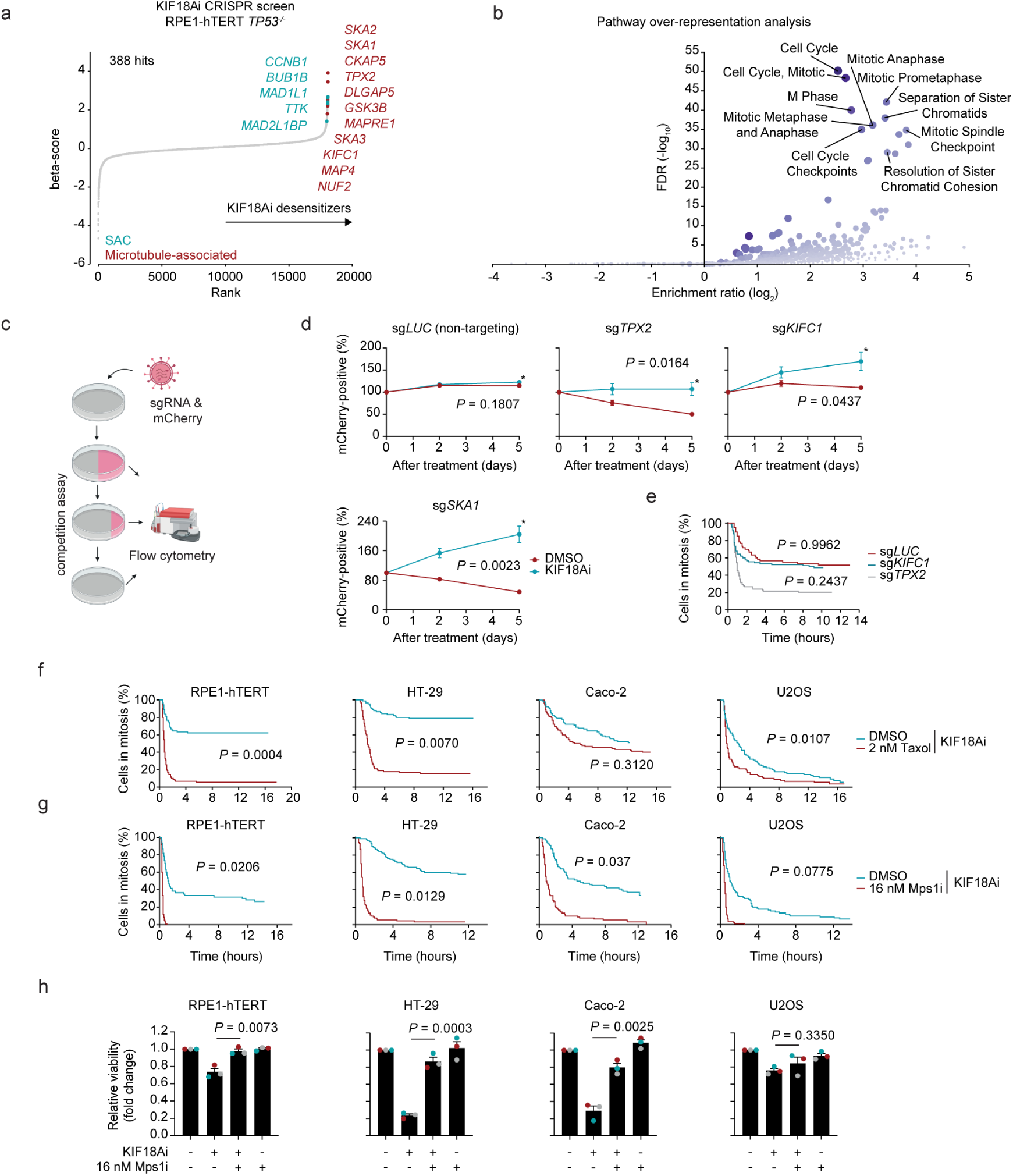
Two convergent mechanisms underlie KIF18Ai resistance. (a) CRISPR knockout viability screen performed in RPE1-hTERT *TP53^-/-^* cells. Cells were treated for 2 weeks with KIF18Ai. The top SAC hits are visualized in blue, while the top microtubule-associated hits are shown in red. Experiment was performed in triplicate. (b) Over-representation analysis of CRISPR screen hits using WebGestalt. (c) Schematic illustrating the setup of the sgRNA-based competition assay. (d) Competition assay results for non-targeting control *luciferase* (LUC) and screens hits *KIFC1*, *TPX2* and *SKA1* in the presence of DMSO or KIF18Ai in RPE1-hTERT *TP53^-/-^*cells. Experiment was performed in triplicate (mean ± s.e.m., two-tailed unpaired t-test on data of day of last measurement). (e) Time in mitosis in the presence of KIF18Ai for cell lines infected with indicated sgRNAs. Experiment was performed in triplicate (one-way ANOVA with Dunnett’s multiple comparisons test comparing sg*LUC*/sg*TPX2* and sg*LUC/*sg*KIFC1*; *n* = 90 cells). (f) Quantification of the time in mitosis of cells treated with KIF18Ai and DMSO, taxol or (g) Mps1i. Experiment was performed in triplicate (Mann-Whitney test; *n* = 90 cells). (h) Cell viability of cells after KIF18Ai and Mps1i treatment. Experiment was performed in triplicate (mean ± s.d., two-tailed unpaired t-test).

Consistent with earlier genetic screens^4, 8^, loss of genes encoding SAC components or microtubule-associated proteins conferred resistance to KIF18Ai (Figure 2a). We validated these findings using a KIF18Ai-dependent competition assay and by measuring mitotic timing following lentiviral expression of mCherry together with sgRNAs targeting *luciferase* (non-targeting control), *KIFC1*, *SKA1*, or *TPX2* (Figure 2c–e). In addition, partial inhibition of the SAC kinase Mps1 using Cpd-5 or stabilization of microtubules with taxol similarly rescued KIF18Ai-induced toxicity (Figure 2f–h). Together, these results indicate that loss of cell viability following KIF18Ai treatment can be alleviated either by overriding the SAC or by restoring microtubule dynamics.

To test whether exposed cancer cells acquired resistance to KIF18Ai through SAC override, we assessed SAC strength using high-dose nocodazole to fully depolymerize microtubules. While nocodazole alone induces a robust mitotic arrest, combining it with a low concentration of Mps1 inhibitor (Mps1i) allows detection of more subtle changes in checkpoint strength^15^. Under these conditions, exposed HT-29 cells spent significantly less time in mitosis than naïve cells, with similar trends observed in Caco-2 and U2OS cells (Figure 3a). However, the average reduction in mitotic duration was much more pronounced in HT-29 cells (42.68 ± 25.30%) than in Caco-2 (15.3 ± 7.59%) or U2OS (11.63 ± 3.54%). These data suggest that SAC weakening is a major resistance mechanism in HT-29 cells but only partially accounts for resistance in Caco-2 and U2OS cells. We therefore conclude that cancer cells may acquire resistance to KIF18Ai by overriding the SAC.

**Figure 3.**
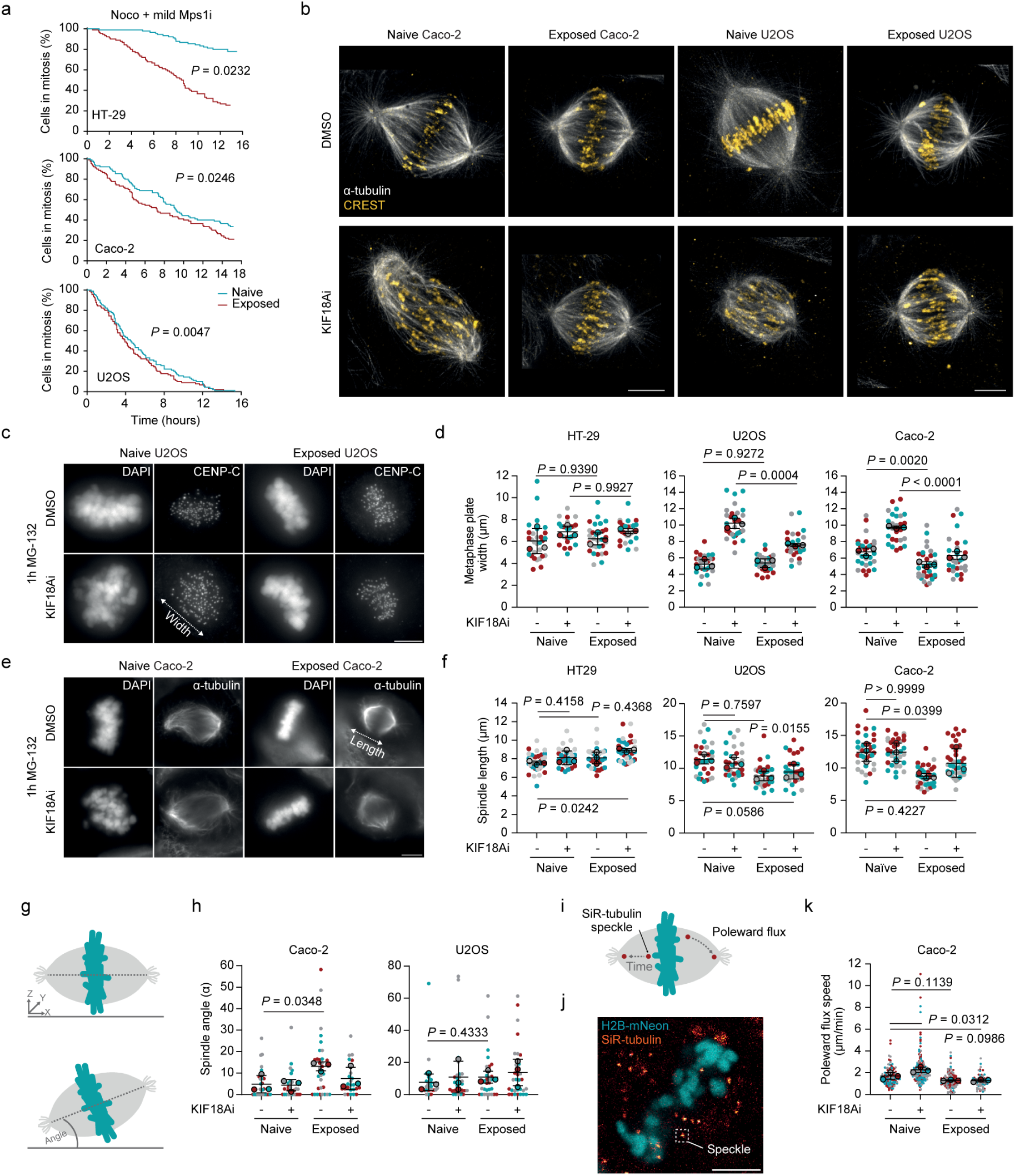
Cancer cells survive KIF18Ai by overriding the SAC or altering the mitotic spindle. (a) Mitotic timings of naïve and exposed cells treated with nocodazole and low Mps1i. Experiment was performed in triplicate (two-way unpaired t-test; *n* = 90 cells). (b) Representative super resolution microscopy images of naïve and exposed Caco-2 and U2OS cells treated with DMSO or KIF18Ai. Images are framed with black borders for presentation purposes. Scale bar = 5 µm. (c) Representative images, and (d) quantification of the metaphase plate width of naïve and exposed cells with and without KIF18Ai. Three independent experiments were performed (mean ± s.d., one-way ANOVA with Šidák’s multiple comparisons test; *n* = 31, 29, 30, 28, 29, 29, 31, 31, 29, 30, 31, 31, respectively). Scale bar = 5 µm. (e) Representative images, and (f) quantification like in d, e, but then for mitotic spindle length. Three independent experiments were performed (mean ± s.d., one-way ANOVA with Dunnett’s multiple comparisons test; *n* = 38, 32, 33, 35, 38, 33, 37, 39, 31, 32, 32 and 33, respectively). Scale bar = 5 µm. (g) Schematic displaying spindle tilt measurement. (h) Quantification of spindle tilt in naïve and exposed cancer cells. Experiment was performed in triplicate (mean ± s.d., two-tailed unpaired t-test; *n* = 31, 29, 32, 24, 30, 29, 30, 30, respectively). (i) Schematic illustrating speckle microscopy concept. (j) Example still of speckle microscopy, and (k) quantification of poleward flux speed. Experiment was performed in triplicate (mean ± s.d., one-way ANOVA with Dunnett’s multiple comparisons test; *n* = 119,176,120 and 77 individual speckles, respectively). Scale bar = 5 µm.

### Spindle microtubule dynamics are rewired in KIF18Ai-resistant cells

Caco-2 and U2OS cells may acquire resistance to KIF18Ai by restoring proper chromosome congression. To examine this more closely, we visualized mitotic spindles and metaphase plates using super-resolution STED microscopy (Figure 3b). This revealed poorly defined kinetochore microtubules and metaphase plates in naïve cells treated with KIF18Ai, whereas exposed cells showed well-defined kinetochore bundles and metaphase plates in both cell lines, irrespective of inhibitor presence. To examine efficiency of chromosome congression more closely, we quantified metaphase plate width by confocal microscopy as a proxy for chromosome misalignment (Figure 3c–d). Following KIF18Ai treatment, naïve Caco-2 and U2OS cells displayed markedly widened metaphase plates, whereas this defect was substantially reduced in exposed cells. In Caco-2 cells, metaphase plate width was reduced by 35% in KIF18Ai-treated exposed cells compared with naïve cells (9.73 ± 0.16 µm versus 6.31 ± 0.46 µm), while U2OS cells showed a 26% reduction (10.24 ± 0.62 µm versus 7.59 ± 0.17 µm). After withdrawal of KIF18Ai, exposed Caco-2 cells exhibited even narrower metaphase plates than naïve cells (23% reduction; 6.75 ± 0.45 µm versus 5.22 ± 0.34 µm), consistent with compensatory adaptation to prolonged KIF18Ai exposure. In contrast, metaphase plate width in HT-29 cells was largely unchanged after KIF18Ai treatment, supporting the conclusion that SAC weakening is the dominant resistance mechanism in this line.

Restored chromosome alignment under KIF18Ai conditions may reflect a re-establishment of baseline microtubule dynamics, a phenomenon previously observed following taxol treatment^8, 12^. To test this idea, we measured mitotic spindle length and spindle tilt relative to the substrates as established proxies for microtubule behavior^16, 17^. Mitotic spindles were significantly longer in untreated naïve cells than in exposed Caco-2 cells (12.40 ± 1.40 µm versus 8.7 ± 0.42 µm) and U2OS cells (11.4 ± 0.70 µm versus 8.83 ± 0.74 µm) (Figure 3e–f), consistent with adaptive changes in microtubule dynamics. In addition, exposed Caco-2 cells showed a significant increase in spindle tilt compared with naïve cells (Figure 3g–h). To directly assess microtubule dynamics, we quantified microtubule poleward flux in metaphase cells using speckle microscopy (Figure 3i–j)^18^, which relies on low concentrations of fluorescent tubulin to generate a speckled-spindle pattern. As reported previously for KIF18A depletion^18^, KIF18Ai increased flux speed in naïve Caco-2 cells (1.70 ± 0.32 µm/min versus 2.26 ± 0.26 µm/min) (Figure 3k). In contrast, the microtubule flux rate in exposed Caco-2 cells did not differ significantly from that in naïve cells (1.30 ± 0.05 µm/min) and remained unchanged following KIF18Ai treatment (1.28 ± 0.12 µm/min), indicating that KIF18A no longer regulates microtubule flux in exposed cells. Together, these findings indicate that dampened microtubule dynamics underlies KIF18Ai resistance in Caco-2 and U2OS cells, rendering KIF18A functionally redundant in resistant cells.

### Adaptation to KIF18Ai confers robust mitotic tolerance rather than dependency

We next asked whether these mitotic adaptations create exploitable vulnerabilities, such as dependence on continued KIF18Ai exposure for proper mitotic progression. To test this, we performed fluorescent live-cell imaging of H2B-mNeon–expressing naïve and exposed cells following KIF18Ai withdrawal and quantified chromosome segregation errors and mitotic timing. Exposed Caco-2, HT-29, and U2OS cells missegregated chromosomes at rates comparable to those of their naïve counterparts (Figure 4a–b), and mitotic timing was unchanged (Figure 4c). These results indicate that exposed cells do not become dependent on KIF18Ai for short-term mitotic progression or survival.

**Figure 4.**
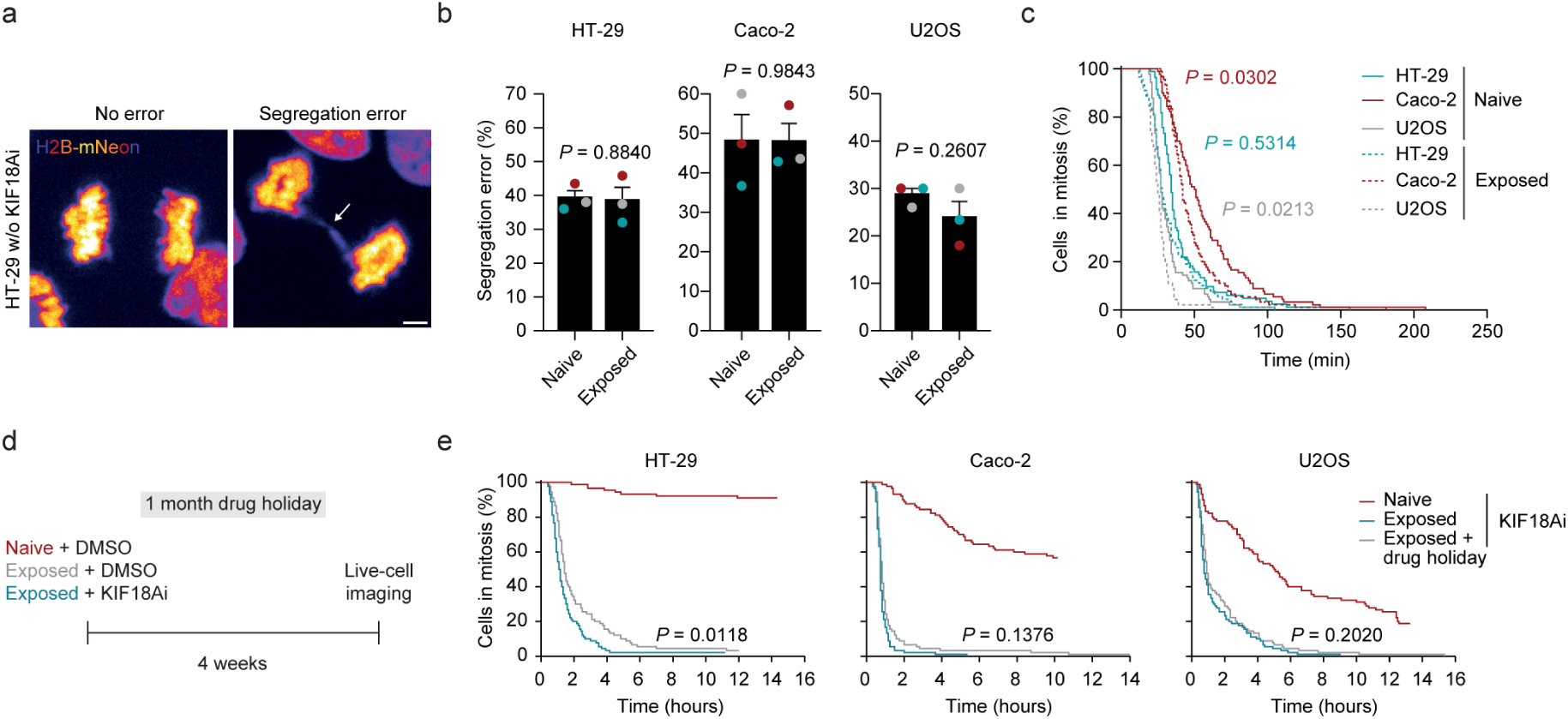
Resistant cancer cells are not KIF18Ai-addicted or resensitized after a drug holiday. (a) Representative stills, and (b) quantification of chromosome segregation errors in naïve and exposed cancer cell lines without KIF18Ai. Three independent experiments were performed (mean ± s.d., two-tailed unpaired t-test; *n* = 90 cells). Scale bar = 20 µm. (c) As in b, but instead mitotic timing was quantified (two-tailed unpaired t-test). (d) Schematic showing the setup of the drug holiday experiment. (e) Quantification of the time in mitosis of naïve, exposed and exposed cells that underwent a drug holiday. Imaging experiments were performed in triplicate (two-tailed unpaired t-test; *n* = 90 cells).

Cancer cells can sometimes lose drug resistance after a prolonged drug holiday if resistance mechanisms impose growth disadvantages^19^. Although we did not observe strong fitness costs in the short term, we asked whether more subtle disadvantages might render KIF18Ai-exposed cells sensitive again to treatment. To test this, we cultured both naïve and exposed cancer cells for four weeks in the absence of KIF18Ai, while maintaining a parallel group of exposed cells under continuous KIF18Ai treatment as a control (Figure 4d). Reintroduction of KIF18Ai immediately before imaging showed that cells that underwent a drug holiday displayed mitotic timing comparable to continuously exposed cells (Figure 4e). These results indicate that resistance to KIF18Ai is stable and that a treatment break does not restore sensitivity.

To examine whether KIF18Ai-exposed cells exhibit increased sensitivity to adjuvant therapies, we performed a small-scale mitotic drug screen (Figure S3b). We focused on exposed Caco-2 cells, as they showed the most pronounced alterations in mitotic spindle morphology (Figure 3f). Thirteen compounds targeting key mitotic processes, including error correction, SAC signaling, microtubule dynamics, and microtubule–kinetochore stability, were tested across ten concentrations each. Unexpectedly, KIF18Ai-exposed Caco-2 cells were not more sensitive to any of the tested compounds compared with naïve cells (Figure S3c). These findings indicate that adaptation to KIF18Ai does not confer increased sensitivity to commonly used mitotic inhibitors, indicating robust tolerance of acquired KIF18Ai-resistant cancer cells to mitotic perturbations.

### Reduced PP2A or APC/C activity sensitizes cells to KIF18Ai

Our results so far show that resistance to KIF18Ai frequently and rapidly arises *in vitro*, highlighting the need to identify tumor contexts with the highest intrinsic sensitivity in order to limit the development of resistance. We therefore asked whether specific genetic alterations could sensitize cancer cells to KIF18Ai and help identify patient populations most likely to benefit from this therapeutic approach. To address this, we examined the results of our CRISPR screen and focused on genes that showed synthetic sickness with KIF18Ai (Figure 5a). In addition to genes whose loss promoted resistance, the screen identified numerous sensitizing interactions involving microtubule-associated genes, including *KIF22*, *CLASP1*, *CLASP2*, *NUMA1*, and multiple dynein subunits, as well as SAC-related genes. These included PP2A-B56 complex members (*PPP2CA*, *PPP2R5C*, *PPP2R5D*, and *PPP2R5E*) and components of the APC/C complex (*UBE2S*, *UBE2C*, and *ANAPC4*).

**Figure 5.**
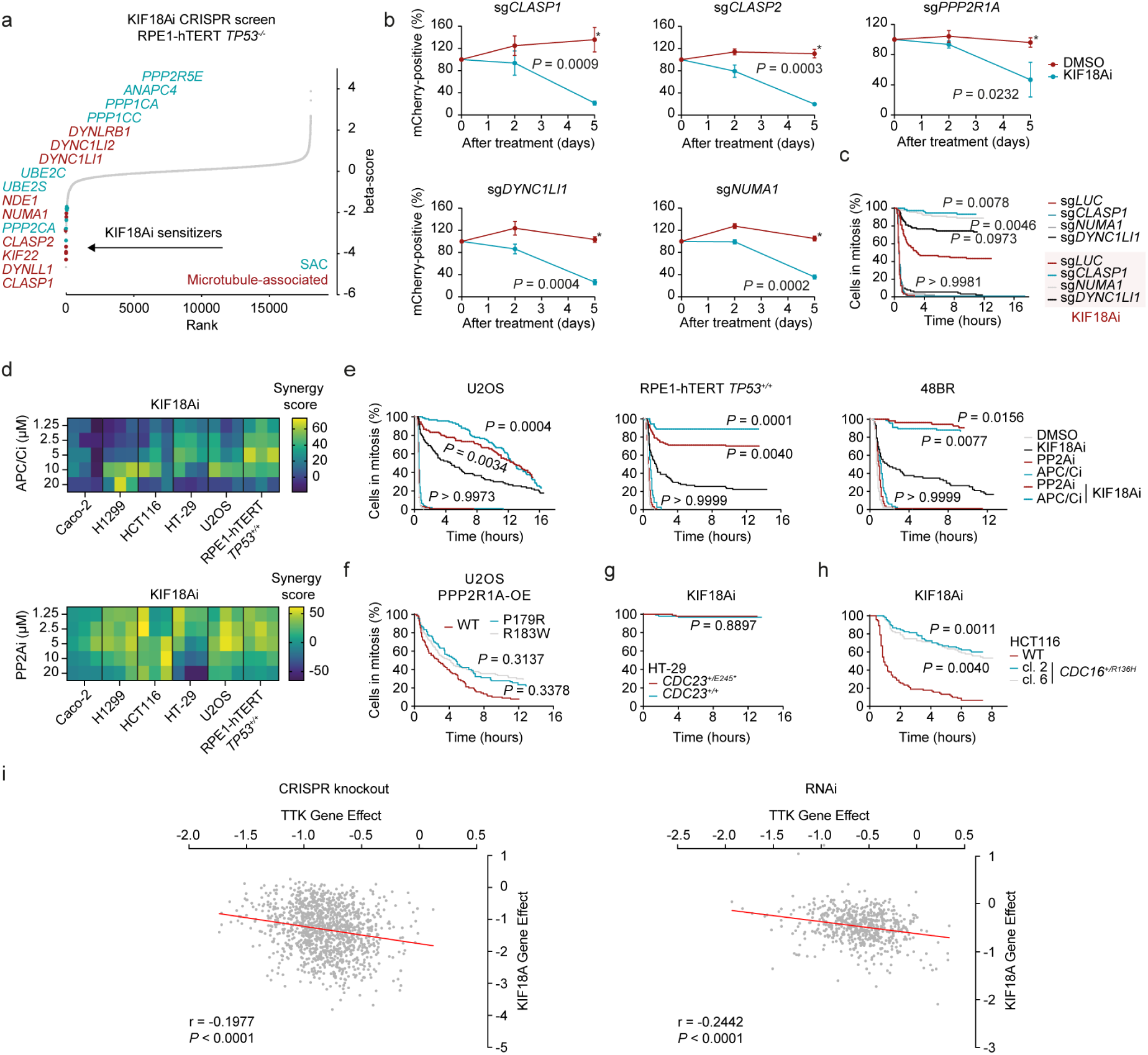
KIF18Ai synergizes with a mitotic exit block via lower PP2A or APC/C activity. (a) CRISPR knockout screen as shown in Figure 2a, but instead the top KIF18Ai sensitizer hits are highlighted. (b) Competition assay in DMSO or KIF18Ai for indicated guides in RPE1-hTERT cells. Experiment was performed in triplicate (mean ± s.e.m., two-tailed unpaired t-test). (c) Mitotic timings of RPE1-hTERT cells after acute depletion using indicated sgRNAs. Experiment was performed in triplicate (one-way ANOVA with Šidák’s multiple comparisons test comparing sg*LUC* with other guides under similar treatment; *n* = 74 cells for sg*CLASP1* + KIF18Ai, 80 cells for sg*TPX2* + KIF18Ai and 90 cells for the others). (d) ZIP model synergy scores^45^ of indicated cell lines after combined treatment with KIF18Ai and APC/Ci or PP2Ai. Experiment was performed in triplicate. (e) Time in mitosis of indicated cell lines after treatment with indicated drugs. Three independent experiments were performed (one-way ANOVA with Šidák’s multiple comparisons test between DMSO-PP2Ai, DMSO-APC/Ci, KIF18Ai-KIF18Ai/PP2Ai or KIF18Ai-KIF18Ai/APC/Ci; *n* = 90 cells, except for 48BR cells (*n* = 85, 77, 83, 72, 52 or 66 cells, respectively). (f-h), Time in mitosis after KIF18Ai of cancer cell lines harboring mutations in PP2A or APC/C subunit genes. Experiments were performed in triplicate (f, h) One-way ANOVA with Dunnett’s multiple comparisons test. (g) Two-tailed unpaired t-test (*n* = 90 cells). (i) Correlation between CRISPR KO (left) and RNAi (right) gene effect scores for KIF18A and TTK across cancer cell lines. Each point represents a cell line. Gene effect scores are from DepMap screens where a more negative values indicate stronger dependency.

The synthetic sickness between KIF18Ai and these factors suggests enhanced SAC activation, either because these proteins function in parallel pathways that promote chromosome alignment, like KIF22, CLASPs or NUMA1^20–22^, or because they directly promote mitotic exit, like members of PP2A-B56 and APC/C complexes^23–25^. We validated five representative hits, including *CLASP1*, *CLASP2*, *DYNC1LI1*, *NUMA1*, and *PPP2R1A*, using competition assays and live-cell imaging to quantify mitotic timing (Figure 5b–c). In addition, combined treatment with KIF18Ai and increasing concentrations of either the PP2A inhibitor LB-100 or the APC/C inhibitor proTAME resulted in a synergistic reduction in cell viability (Figure 5d). Consistently, treatment of multiple cell lines with moderate doses of PP2A inhibitor (2.5 µM) or APC/C inhibitor (5 µM) strongly exacerbated the KIF18Ai-induced mitotic arrest, whereas either inhibitor alone had minimal effects (Figure 5e). Together, these findings indicate that reduced activity of specific microtubule-associated or SAC-related proteins synergizes with KIF18Ai.

Having established that reduced PP2A or APC/C activity enhances KIF18Ai sensitivity, we next asked whether clinically relevant mutations in these pathways could modulate KIF18Ai toxicity. Mutations in the PP2A regulatory subunit *PPP2R1A* occur in several cancer types, affecting approximately 0.3% of tumors and typically presenting as heterozygous missense mutations^26^. These mutations have been linked to activation of oncogenic signaling pathways. To test whether such alterations increase KIF18Ai sensitivity, we overexpressed wild-type PPP2R1A or two common dominant-negative PPP2R1A mutants (P179R and R183W)^27, 28^ in U2OS cells (Figure S3d). This resulted in only a mild and non-significant increase in mitotic duration following KIF18Ai treatment (Figure 5f).

In addition to PP2A, mutations in genes encoding for core APC/C components such as *CDC16*, *CDC23*, and *CDC27* have been reported in tumors and are thought to limit excessive chromosomal instability by prolonging mitosis^29^. To assess whether such mutations enhance responsiveness to KIF18Ai, we analyzed multiple cell lines, including HT-29 cells carrying either a pre-existing *CDC23^+/E2^*^45^*\** mutation or a corrected *CDC23^+/+^*allele, as well as two Mps1i–adapted HCT116 clones harboring a *CDC16^+/R136H^*mutation^29^. KIF18Ai treatment did not reveal differences in mitotic arrest between mutated and corrected HT-29 cells (Figure 5g). In contrast, both *CDC16^+/R136H^* HCT116 clones exhibited significantly prolonged mitotic arrests compared with the parental line upon KIF18Ai treatment (Figure 5h), suggesting Mps1i resistance can create vulnerabilities that render cancer cells more susceptible to KIF18Ai. To evaluate the generalizability of this observation, we plotted the sensitivity of more than 1,100 cancer cell lines to loss of *TTK* (encoding Mps1) versus KIF18A using data from the Cancer Dependency Map^30^. This analysis revealed a striking anticorrelation between the two dependencies across both CRISPR knockout and RNAi experiments (Figure 5i), further indicating that resistance to loss of Mps1 activity increases sensitivity to KIF18A inactivation. Overall, while not all clinically observed PP2A or APC/C mutations substantially alter KIF18Ai sensitivity, our data demonstrate that mitosis-prolonging alterations acquired during Mps1i resistance can make cells particularly sensitive to KIF18Ai (Figure 6).

**Figure 6.**
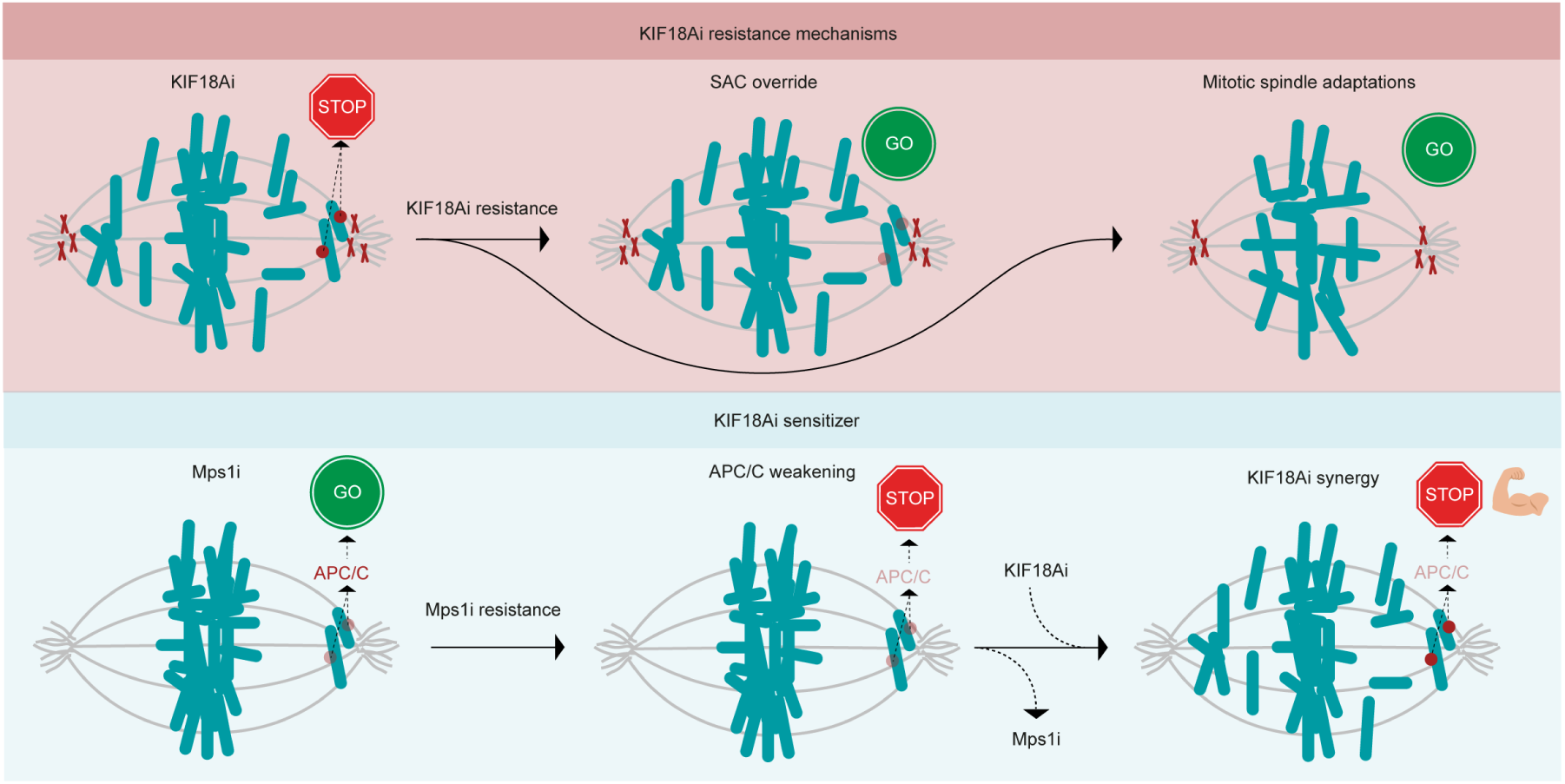
Distinct resistance and sensitization pathways link SAC signaling, APC/C activity, and KIF18A inhibition. Top: KIF18Ai resistance mechanisms. Acute KIF18A inhibition blocks mitotic progression (STOP). Cells that acquire resistance bypass the spindle assembly checkpoint (SAC override), allowing mitotic exit despite persistent chromosome alignment problems (GO). Continued adaptation may also lead to mitotic spindle remodeling that supports division under KIF18Ai. Bottom: KIF18Ai sensitization via MPS1i resistance. Inhibition of Mps1i promotes mitotic progression through APC/C activation (GO). Prolonged Mps1i treatment selects for resistance associated with partial APC/C weakening, restoring normal mitotic duration (STOP). In this context, KIF18Ai sensitivity increases (stronger STOP).

## Discussion

Loss of KIF18A activity is highly lethal in specific cancer cell contexts^5–7, 10^. Here, we show that these same cells can rapidly acquire resistance to KIF18A inhibition. Given that KIF18A inhibitors are currently being evaluated in clinical trials, our findings suggest that resistance is also likely to emerge *in vivo*. We find that resistance arises through two broad strategies (Figure 6). Cells either restore chromosome congression by reducing their reliance on KIF18A for chromosome alignment, or they weaken the spindle assembly checkpoint (SAC) to escape the otherwise toxic mitotic arrest. Together, these results underscore the diverse adaptive mechanisms cancer cells can use to survive KIF18Ai-induced stress and point to multiple pathways that may shape therapeutic response. Consistent with this, initial sensitivity to KIF18Ai has also been linked to the induction of severe chromosome misalignment^5, 7^ and the maintenance of a strong mitotic arrest^8^. Therefore, when either of these conditions is not or only partially met, cells are able to tolerate and survive KIF18A inhibition.

The molecular basis of resistance to KIF18Ai in our cell lines remains incompletely defined and is likely complex. Our CRISPR knockout screen identified hundreds of genes whose loss either sensitized cells to KIF18Ai or promoted resistance, suggesting that resistance can arise through diverse genetic or regulatory changes. While individual genes may contribute, overall sensitivity may reflect the combined effects of multiple pathways. Mitotic spindle regulation is highly redundant and depends on a finely tuned balance of forces and components^31, 32^, making it plausible that the microtubule-stabilizing role of KIF18A becomes dispensable in some resistant cells. Such adaptation could arise, for example, through altered activity of microtubule nucleators such as TPX2^33^ or CKAP5^34^. Likewise, cells that override the SAC to escape KIF18Ai-induced arrest may do so by reducing SAC signaling through diminished Mps1, MAD1, or MAD2 activity, or by increasing the activity of mitotic exit regulators such as PP2A or the APC/C, as previously noted^8^.

Our findings that resistance to KIF18Ai frequently emerges during treatment emphasizes the need to develop strategies to prevent or overcome it. We observed strong synergy between KIF18Ai and pharmacological inhibition of PP2A or the APC/C, consistent with prior observations following APC/C subunit depletion^35^, combined APC/C inhibition and microtubule poisons, or reduced APC/C activity in cancer cells^8^. These results raise the possibility that combining KIF18Ai with PP2A or APC/C inhibitors could enhance therapeutic efficacy. However, this synergy was not restricted to cancer cells and was also observed in non-transformed cells such as 48BR and RPE1-hTERT, suggesting that such combinations may carry a substantial risk of toxicity. Similarly, combining KIF18A inhibitors with other anti-mitotic drugs requires caution, as both our data and previous studies show that Mps1 inhibition or a low-dose of taxol can suppress the cytotoxic effects of KIF18Ai^8, 12^.

Notably, our results suggest that resistance to loss of Mps1 activity, for example in cells adapted to Mps1i through a R136H mutation in the APC/C subunit CDC16^36^, sensitizes cells to KIF18Ai. Prolonging mitosis through reduced APC/C activity seems to be a common adaptation to Mps1i treatment^37, 38^ and since Mps1 inhibitors are currently under clinical evaluation, these findings point to KIF18A inhibition as a potential vulnerability in a subset of Mps1i-resistant cancers. Interestingly, a recent report shows that low levels of the main APC/C activator, CDC20, reduce sensitivity to Mps1 inhibition in cancer cells, suggesting that CDC20 levels may in turn account for increased sensitivity to KIF18A inhibition^39^. Numerous naturally occurring mutations in APC/C subunits have been identified in tumors^29^. Although we did not observe enhanced KIF18Ai sensitivity in CDC16-mutated HT-29 cells, systematic evaluation of additional APC/C mutations will be important. The synergy between KIF18Ai and reduced APC/C activity may be explained by reciprocal regulation between the SAC and the APC/C. APC/C inhibition triggers a SAC-dependent mitotic arrest^40, 41^ and the SAC and APC/C mutually restrain one another through feedback mechanisms^41, 42^. Concurrent inhibition of both pathways may therefore disrupt this feedback loop and strongly amplify the mitotic arrest.

Despite a substantial reduction in spindle size and dampening of microtubule dynamics in some KIF18Ai-resistant cells, we did not identify new vulnerabilities using a panel of mitotic drugs. This was unexpected and suggests that resistant cells retain broad sensitivity to mitotic perturbations. It remains possible that more targeted disruption of microtubule dynamics, such as further spindle shortening, could expose context-specific vulnerabilities. For example, genetic perturbations that confer resistance to KIF18Ai in naïve cells may instead induce mitotic defects once cells have adapted. Identifying such context-dependent vulnerabilities will be an important direction for future studies.

Finally, we find that although the mitotic machinery in resistant cells adapts to function in the absence of KIF18A, mitotic progression stays at near-normal levels once KIF18A inhibition is removed. At first glance, this is surprising, as SAC weakening or altered microtubule dynamics might be expected to compromise chromosome segregation^43^ or prolong mitosis^11^. However, both a prior study^8^ and our data indicate that only modest SAC weakening^44^ is sufficient to bypass the KIF18Ai-induced arrest, while still allowing accurate cell division after drug removal. Consistent with this, we have shown that low-dose Mps1 inhibition alone does not impair viability but rescues survival when combined with KIF18Ai. Similarly, exposed cells with altered spindle architecture divide normally after KIF18Ai withdrawal, likely because spindle dynamics have been rewired but remain within a functional range. A comparable phenomenon was observed in our CRISPR screen and competition assays following KIFC1 knockout, where cells tolerate spindle alterations under untreated conditions yet become resistant to KIF18A inhibition^8^.

In summary, our study identifies key determinants of sensitivity and resistance to KIF18A inhibition, providing new insight into mitotic adaptation and highlighting both the promises and the challenges of targeting mitosis for cancer therapy.

## Methods

### Cell lines

Caco-2 and HT-29 (a kind gift from the Barnhoorn lab), MDA-MB-231 (a kind gift from the van der Maarel lab), H1299, Hela, U2OS and RPE1-hTERT TP53WT (a kind gift from the van Attikum lab) and RPE1-hTERT iCas9 TP53/Puro double knockout (dKO)^46^ were cultured at 37 °C in at 5% CO2 in DMEM GlutaMAX (Thermo Fisher Scientific) supplemented with 1% penicillin–streptomycin (Sigma-Aldrich) and 10% FBS (Gibco) (see Supplemental Table 1). HT-29 CDC23^+/E245*^ and HT-29 CDC23^+/+^ cell lines were a kind gift from the Swanton lab. HCT116 CDC16^R136H^ clones were a kind gift from the Medema lab. H2B-mNeon-expressing cells were generated by infecting cells with a lentivirus encoding for H2B-mNeon and zeocin^47^ (see Supplemental Table 3 for a list of plasmids). Selection using 20 ug/ml Zeocin (Invivogen) was performed for 7 days. PPP2R1A overexpression cell lines (PPP2R1A overexpression plasmids^48^ were a kind gift from Veerle Janssens) were generated by infecting cells with lentivirus.

### Immunofluorescence

Cells were split at 40% confluency in a 24 well plate on 12-mm round glass coverslips (VWR). The next day, cells were treated for 2 hours with KIF18Ai. In case of kinetochore stainings, pre-extraction was performed in 0.2% of triton in PHEM buffer, followed by fixation in 4% paraformaldehyde (Sigma-Aldrich). For KIF18A staining, cells were fixed in −20 °C 100% methanol (J.T. Baker) for 15 min. Cells were washed three times with PBS, permeabilized in 0.5% Triton-X (Merck)/PBS for 10 min at RT and blocked for 10 min in 3% BSA (Sigma)/PBS at RT. Next, coverslips were incubated for 2-18 hours with primary antibodies (see Supplemental Table 5 for a list of antibodies), followed by 2h of secondary antibodies in 3% BSA/PBS with 4x PBS washes in between. After washing 3x for 5 min with PBS, coverslips were incubated for 5 min with DAPI, washed with PBS and mounted using ProLong Gold Antifade (Invitrogen) on a glass slide. Images were acquired on a Zeiss AxioImager M2 widefield fluorescence microscope equipped with a 63x PLAN APO (1.4 NA) oil-immersion objectives (Zeiss) and an HXP 120 metal-halide lamp used for excitation. For metaphase plate width and spindle tilt experiments, every 0.5 µm images were taken in Z for a total of 8 um. For mitotic spindle length imaging experiments, images were acquired in a single plane. For metaphase plate width and mitotic spindle length experiments, mitotic cells with severe spindle tilt were excluded from the analysis. Images were recorded using ZEN 2012 (Blue edition, v.1.1.0.0) and analysed using Fiji.

Metaphase plate width was measured manually using the line tool by drawing a line perpendicular to the metaphase plate at its widest point, spanning between the two most distant kinetochores as marked by CENP-C. Mitotic spindle length was measured by manually drawing a line between the two microtubule-organizing centers as marked by α-tubulin. Spindle tilt was measured manually by determining the center of both microtubule-organizing centers in Z and XY as marked by α-tubulin and calculating the angle using the formula arctan(ΔZ/ΔXY).

### Metaphase spreads

Cells were split to 30% confluency on 12-mm coverslips in 24 well plates. The next day, cells were incubated for 6 hours in 500 ul culture medium containing 3.3 uM nocodazole, followed by slow addition of 500 ul of 1:7 PBS:MQ. Cells were washed twice with the hypotonic solution and incubated at 37 °C for 30 min. Next, fixation was performed by slowly adding 500 ul of 3:1 methanol:acetic acid to the cells, followed by two washes with the fixation solution. Cells were incubated at room temperature for 10 min. Slides were subsequently air-dried to allow for cell swelling and were stained following the immunofluorescence protocol using CENP-C or CENP-A antibodies (For a list of antibodies see Supplemental Table 5).

### Generation of KIF18Ai exposed cancer cell lines

Cancer cell lines were continuously treated for at least 6 weeks with DMSO or 250 nM of the KIF18Ai sovilnesib (MedChemExpress). Medium was refreshed with new inhibitor whenever the DMSO-treated cells reached confluency and were split, which was approximately every 3-4 days. Cells were maintainedon treatment after the 6 weeks of treatment for at least six weeks.

### Live-cell imaging

Cells were split to 40% confluency in a 24 or 48 wells plate 1 day before imaging. Drugs (3.3 µM nocodazole (Sigma-Aldrich), Mps1i (Cpd-5; Key Organics), taxol (a kind gift from the Kops lab)) were added right before imaging on a Leica AF6000 Inverted using a HC PL FL 10x (0.3 NA) dry objective and brightfield settings in a climate chamber at 37 °C and 5% CO_2_. Pictures were taken overnight every 5 minutes. For acute sgRNA live-cell imaging experiments (see Supplemental Table 2 for a list of gRNAs), cells were infected with lentivirus as described previously and treated with 100 ng/mL doxycycline (Sigma-Aldrich) to induce Cas9 expression. 3-4 days after infection, cells were split to 30% confluency in a 24 well plate and imaged a day later. Images were taken using RFP settings (BP 545-551 excitation and BP 570-640 emission filter). Analyses were manually performed in ImageJ version 1.54f. The time of mitotic entry and exit was defined as the first frame of cell rounding and anaphase entry/cell flattening, respectively. We only quantified mitotic timings of cells entering mitosis during the few first hours of imaging.

Long-term live-cell imaging was mostly performed as described previously^49^. Briefly, the Lattice Lightsheet 7 system (Carl Zeiss, Germany) was used for live-cell imaging of Caco-2 cells expressing H2B-mNeon. The system was equipped with an illumination objective lens (13.3×/0.4, at a 30° angle to the coverslip) with a static phase element and a detection objective lens (44.83×/1.0, at a 60° angle to the coverslip) with an Alvarez manipulator. Images were acquired using ZEN 2.6 (Blue edition; Zeiss). Automatic water immersion was applied from the motorized dispenser at intervals of 20 or 30 min. Immediately after sample mounting, four steps of the “create immersion” auto-immersion option were applied. The sample was illuminated with a 488 nm diode laser (power output 10 mW), with laser power set to 1%. The detection module consisted of a Hamamatsu ORCA-Fusion sCMOS camera with the exposure time set to 15 ms. The LBF 405/488/561/642 emission filter was used, blocking emission detection outside the 420–770 nm range. During imaging, cells were maintainedat 37 °C and 5% CO₂ in a Zeiss stage incubation chamber system (Zeiss). The width of the imaging area in the x dimension was set to 2 mm, with a 0.4 µm interval size. The time between consecutive frames was set to 2 min. The total imaging duration was 4 days and was occasionally interrupted by air bubbles, which caused a loss of intensity in part or all of the imaging area. The light sheet length, also referred to as the field of view or illumination width, was 30 µm, while its thickness was set to 1000 nm. The parameters “Focus sheet,” “Focus waist,” and “Aberration control” were manually fine-tuned before each imaging session, with ranges of −0.200 to −0.230, 50 to 60, and 170 to 185, respectively. Alternatively, a Dragonfly spinning disk confocal microscope system (Andor Technology, Belfast, UK) equipped with a 20×/0.75 NA PL APO air objective (Nikon, Tokyo, Japan) and a Sona 4.2B-6 back-illuminated sCMOS camera (acquisition mode set to high speed; Andor Technology) was used for long-term live-cell imaging. Images were acquired using Fusion 2.4 software. During imaging, cells were maintained at 37 °C and 5% CO₂in a heating chamber (Okolab, Pozzuoli, NA, Italy). For excitation of mNeon, the 488 nm laser line with corresponding green filter was used. Image acquisition was performed every 2 µm with 8 focal planes and a time interval of 5 min, with an xy resolution of 0.6 µm. The total imaging duration was 4 days.

### Viability assays

Cells were split at 5% confluency in a Nunc 96 MicroWell white bottom plate or regular cell culture 96 well plates and treated with indicated drugs. After the indicated time periods, cell viability was measured using MTT (Life Technologies) or the CellTiter-Glo 2.0 kit (Promega) according to the manufacturer’s instructions. For the Caco-2 mini-drug screen, we used the following inhibitors from MedChemExpress, unless indicated otherwise; nocodazole, taxol, CENPEi (GSK-923295), KIF11i (monastrol), KIF15i (Kif15-IN-1), MCAKa (UMK57), MKLP2i (paprotrain), Aurkai (Alisertib; TargetMol), Aurkbi (ZM-447439;TargetMol), Mps1i (Cpd-5), PLK1i (GSK461364), PLK4i (Centrinone) and APC/Ci (proTAME) (also see Supplemental Table 6).

### *KIF18A* sequencing

RNA was extracted using the RNeasy kit (Qiagen) following the instructions of the manufacturer including an on-column DNase treatment. Reverse transcription was performed using TaqMan Reverse Transcription Reagents (Thermo Fisher Scientific). The *KIF18A* gene was amplified, but because the full gene could not be sequenced in a single reaction, 5 primers were used to Sanger sequence overlapping regions covering the entire gene (see Supplemental Table 4 for primers).

### Western blotting

Total cell lysates were harvested by scraping cells in Laemmli-SDS sample buffer. Cell lysates were boiled for 10 min at 95°C. Proteins were separated on Criterion™ XT Bis-Tris 4–12% Protein Gels (Biorad) in MOPS Running Buffer (Thermo Fisher). Then, blotted onto PVDF membranes (IPFL00010, EMD Millipore) in Tris/glycine blotting buffer (0.025 M Tris, 0.192 M glycine) with 20% methanol. Membranes were blocked with commercial blocking buffer (Rockland) for 1 h at room temperature. Membranes were then probed with indicated antibodies in commercial blocking buffer (For antibodies see Supplemental Table 5). Proteins were stained with fluorochrome-conjugated secondary antibodies and were detected on an Odyssey CLx system and Image Studio software (Li-Cor).

### Flow cytometry

DNA content was measured as described before^47^. In short, nuclei were harvested using a nuclear isolation buffer and stained using Hoechst 33342. Flow cytometry was performed on a BD LSRFortessa (BD Biosciences) using a violet laser and 525/550 emission filter. Data was visualized using FlowJo.

### CRISPR screen

RPE1-hTERT iCas9 TP53/Puro dKO cells were infected with the Toronto human KnockOut (TKOv3) pooled library^50^ (a kind gift from the Moffat lab; Addgene #125517) at an MOI of <0.35 at >100x coverage using 8 µg/ml polybrene for 1 day and 100 ng/ml doxycycline for 3 days. Selection using 3 µg/ml puromycin was performed 1 day after infection for 2 days. T=0 was harvested 4 days after infection and a 15 day KIF18Ai treatment was started 5 days after infection. Cell pellets were stored at −20 °C until genomic DNA was isolated using the Wizard Genomic DNA purification kit (Promega) using the instructions of the Moffat lab under header 3.5.2 (see Addgene #125517). sgRNA sequences were amplified using a two-step PCR as described previously by the Moffat lab (see Addgene #125517) (For primers see Supplemental Table 4). After the first PCR, the product was purified using 0.6x volume Ampure XP reagent beads (Beckman Coultier) and at 1.8x volume after the second PCR. Fragments were sequenced by Macrogen on an Illumina NovaSeq at 2×150 bp. Reads were trimmed to 1×50 bp and the analysis was performed using the MAGeCK2 algorithm. Over-representation analysis was performed by taking all significant (*P <* ^51^⍰.

### Competition assay

Cells were infected (<60% transduced cells) with lentivirus encoding for indicated sgRNAs (table 1) and mCherry (lentiviral transfer plasmid pLenti-guide-OSS-mCherry^52^. Percentage of mCherry-positive cells were determined at indicated times using an ACEA Novocyte flow cytometer using a 561 nm laser and a 615/20 detector while gating for single cells using the area of the forward and side-scatter, the height of the side-scatter versus the area of the side-scatter and the height of the forward scatter versus area of the forward scatter. ∼2000 cells were measured for each condition.

### Super-resolution microscopy

Super-resolution microscopy was performed as described previously^49, 53^. Briefly, parental WT Caco2 and U2OS cells were washed with cell extraction buffer (CEB) and fixed for 10 min at room temperature with 3.2% paraformaldehyde (PFA) and 0.1% glutaraldehyde (GA) in PEM buffer (0.1 M PIPES, 1 mM MgCl₂·6H₂O, 1 mM EGTA, 0.5% Triton X-100). Following fixation, residual aldehydes were quenched by incubation in freshly prepared 0.1% sodium borohydride in PBS for 7 min, followed by incubation in 100 mM glycine in PBS for 10 min at room temperature. Cells were blocked and permeabilized in blocking buffer containing 2% normal goat serum and 0.5% Triton X-100 for 2 h at room temperature. Primary and secondary antibody incubations were performed as described previously for immunofluorescence experiments. Primary antibodies were: rat anti-tubulin (1:500, MA1-80017, Invitrogen) and human anti-centromere protein (1:500, 15-234, Antibodies Incorporated). Secondary antibodies were: donkey anti-rat IgG Alexa Fluor 594 (1:300, ab150156, Abcam) and donkey anti-human IgG DyLight 488 (1:1000, ab102424, Abcam). Super-resolution imaging was performed using an Expert Line easy3D STED microscope system (Abberior Instruments) equipped with a 100×/1.4 NA oil immersion objective (UPLSAPO100x, Olympus) and avalanche photodiode (APD) detectors. Excitation was performed using the 488 nm (set to 10% relative power) and 561 nm (set to 30% relative power) laser lines, with a 775 nm depletion laser (set to 20% relative power) for STED imaging of red channel. For fixed-cell imaging, the xy pixel size was set to 20 nm, and z-stacks consisting of 12 optical sections were acquired with a step size of 200 nm. Dwell time was set to 10 µs, and no additional line accumulations were used. Default detection filters were used.

### Speckle microscopy

Speckle microscopy was performed as described previously^18^, with few modifications. In short, Caco-2 H2B-mNeon-expressing cells cultured on glass-bottom dishes were labeled with 0.5–1 nM SiR-tubulin (Spirochrome AG) and incubated for 15 min prior to imaging. Live-cell confocal microscopy was then performed using an Expert Line easy3D STED microscope (Abberior Instruments) fitted with a 100×/1.4 NA UPLSAPO100x oil immersion objective (Olympus) and avalanche photodiode detectors. Image acquisition was controlled using Imspector software. Throughout imaging, cells were maintainedat 37 °C in a humidified atmosphere containing 5% CO₂ using a stage-top incubation system (Okolab). Excitation was achieved using 485 nm (1% relative power) and 640 nm (80% relative power) laser lines. Time-lapse sequences were recorded at a single focal plane with intervals of 5 or 10 s. Dwell time was set to 10 µs with 3 or 4 line accumulations and a pixel size of 90 nm. Default detection filters were used.

Microtubule speckle dynamics were quantified using kymograph-based analysis in Fiji (ImageJ). Time-lapse image sequences were first rotated so that the main spindle axis was aligned horizontally, without interpolation, to preserve pixel integrity. A narrow rectangular region of interest (ROI; 10–50 pixels in thickness) was drawn along the spindle axis, spanning the length of the spindle. Kymographs were generated using the Reslice function (Image → Stacks → Reslice), with an output spacing of 1 pixel and without applying any projection along the time axis, ensuring preservation of temporal information. In the resulting kymographs, the horizontal axis corresponds to spatial position along the spindle axis, and the vertical axis corresponds to time (frames). Image calibration was applied such that spatial measurements were converted using the pixel size of 0.090 µm/pixel, and temporal measurements were converted using the acquisition interval of 5 s per frame. Individual microtubule speckles were manually tracked by drawing straight lines along their trajectories in the kymographs. Speckle displacement (Δx) and elapsed time (Δt) were extracted from the line coordinates, and speckle velocities were calculated as Δx/Δt and expressed in µm/min. Speckle duration was estimated independently from spatial displacement by measuring the total time interval over which each speckle remained visible in the kymograph and converting this value to seconds using the frame interval. Only clearly discernible, continuous speckle trajectories were included in the analysis. Cells with less than 5 clearly observable speckles were excluded from the analysis.

### Statistics

Statistical analyses were performed in GraphPad Prism 10.2.3. To determine statistical significance of mitotic timings, the average time spent in mitosis was used. The mean values were normalized to controls in case of SAC strength assays due to variability between independent experiments.

## Acknowledgments

We thank the Barnhoorn lab for providing the Caco-2 and HT-29 cell lines, and the van der Maarel lab for the MDA-MB-231 cell line. We are grateful to the Swanton lab for the HT-29 *CDC23^+/E2^*^45^*\** and HT-29 *CDC23^+/+^*cell lines, the Medema lab for the HCT116 *CDC16^R136H^* clones and the de Wind lab for HCT116 cells. We thank Veerle Janssens for providing the PPP2R1A overexpression plasmids and the Kops lab for the generous gift of taxol. We also acknowledge the Kops lab for the H2B-mNeon construct used in this study.

## Author contribution

SJK conceived and designed the study, performed experiments unless stated otherwise, analyzed and interpreted the data unless stated otherwise, and wrote the manuscript. KV provided conceptual input, assisted with super-resolution microscopy, long-term imaging and speckle microscopy experiments and performed speckle microscopy analysis. MvG provided support for the CRISPR/Cas9 screens and generated CRISPR screen virus. HvA supervised MvG. IT supervised KV. MSL supervised the project and wrote the manuscript.

---

**Supplemental Table 1:** Cell lines.

| **Cell line (ATCC number)** | **Source** |
| --- | --- |
| Caco-2 (HTB-37) | Barnhoorn lab |
| H1299 dox-inducible shBRCA2 (CRL-5803) | Noordermeer lab |
| HCT116 (CCL-247) | De Wind lab |
| HCT116 CDC16 R136H (CCL-247) | Medema lab |
| Hela (CRM-CCL-2) | Van Attikum lab |
| HT-29 (HTB-38) | Barnhoorn lab |
| HT-29 CDC23 +/+ (HTB-38) | Swanton lab |
| HT-29 CDC23 +/-E245* (HTB-38) | Swanton lab |
| MDA-MB-231 (CRM-HTB-26) | Van der Maarel lab |
| RPE1 TetOn iCas9 Puro/p53-dKO (hereafter RPE1-hTERT TP53 KO) (CRL-4000) | Wolthuis lab |
| RPE1-hTERT (hereafter RPE1-hTERT TP53WT) (CRL-4000) | Van Attikum lab |
| U2OS (HTB-96) | Van Attikum lab |

**Supplemental Table 2:**
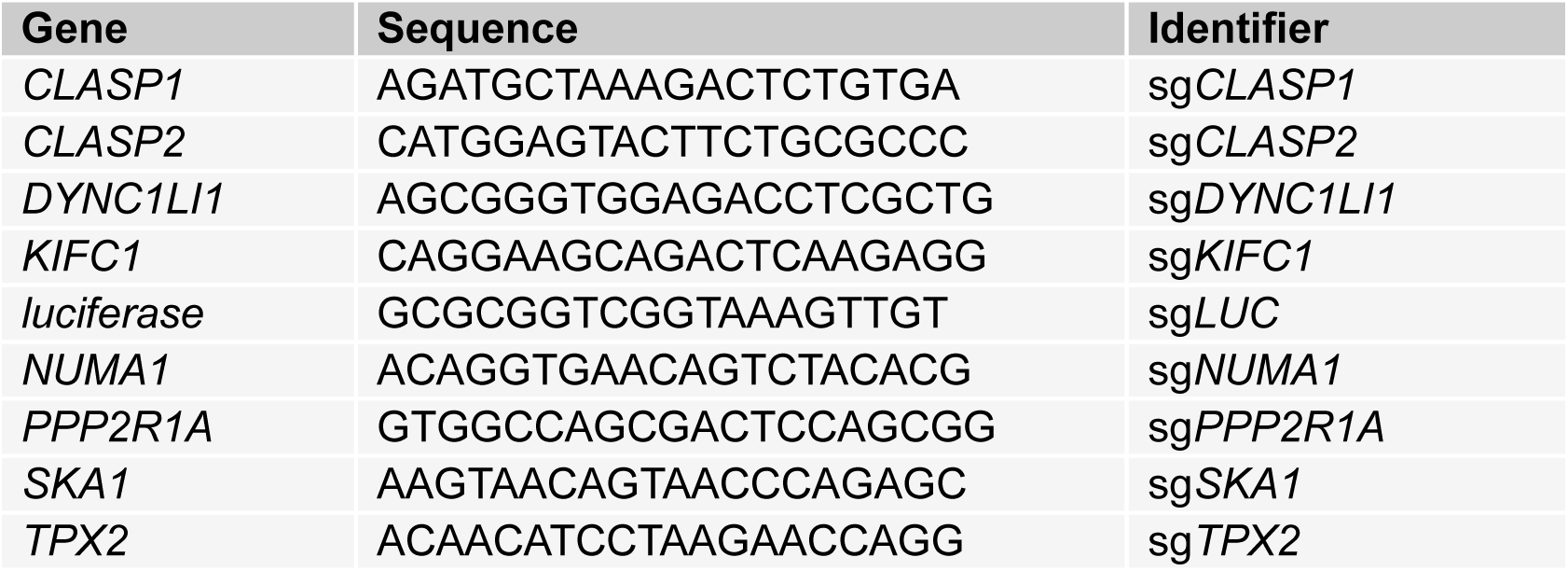
sgRNAs.

**Supplemental Table 3:** Plasmids.

| Plasmid | Origin |
| --- | --- |
| pCMV-VSV.G | Addgene (#8454) |
| pLenti6.4_PPP2R1A_P179R_3xFLAG_Blast | Janssens lab |
| pLenti6.4_PPP2R1A_R183W_3xFLAG_Blast | Janssens lab |
| pLenti6.4_PPP2R1A_WT_3xFLAG_Blast | Janssens lab |
| pLenti-guide-OSS-mCherry | Luijsterburg lab |
| pLV-H2B-Neon-ires-Zeocin | Kops lab |
| pMDLg-pRRE | Addgene (#12251) |
| pRSV-REV | Addgene (#12253) |
| Toronto KnockOut (TKO) CRISPR Library - Version 3 | Addgene (#125517) |

**Supplemental Table 4:**
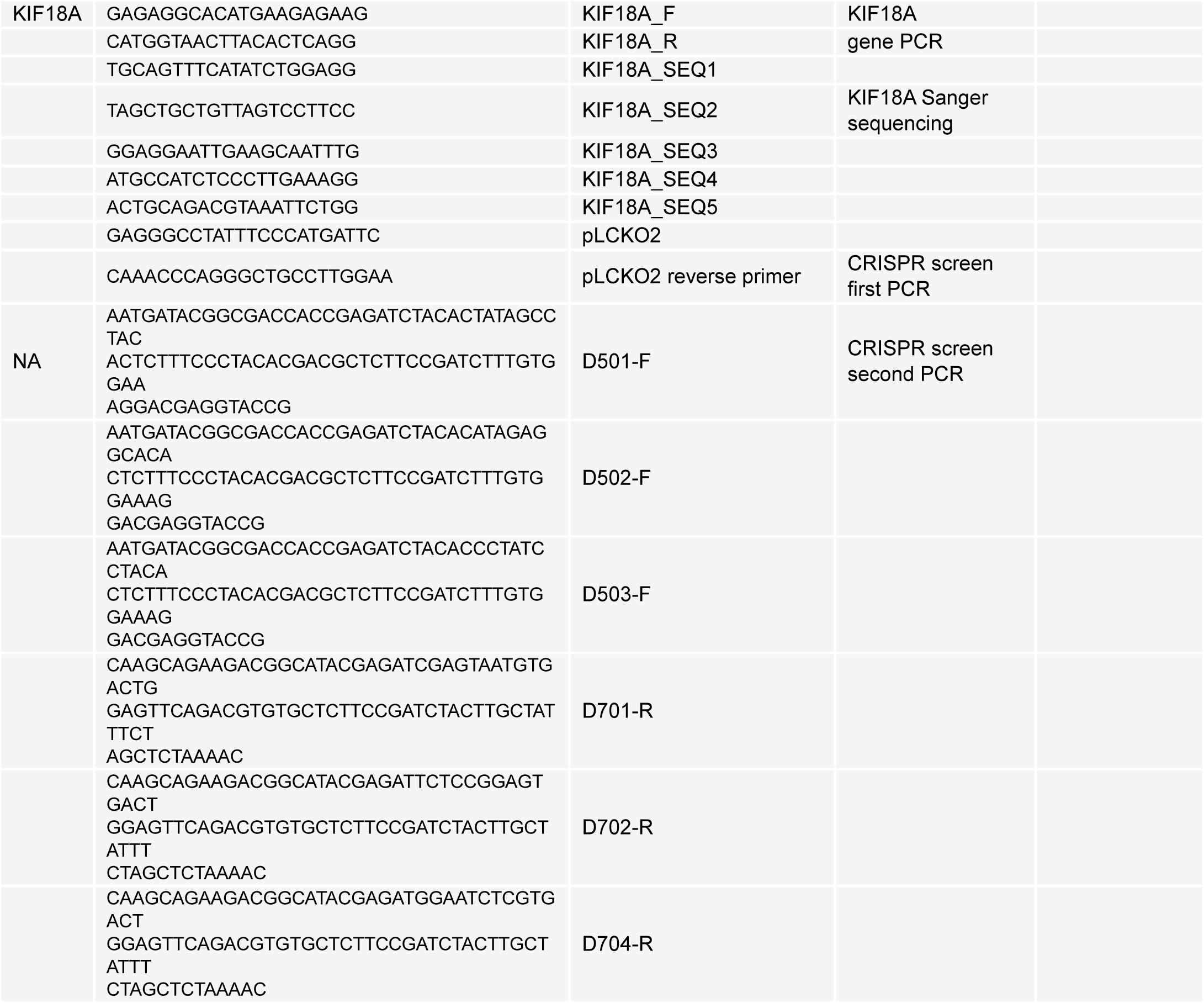
Primers.

**Supplemental Table 5:** Antibodies.

| **Antibody** | **Host** | **Company (reference)** | **Use** | **Identifier** |
| --- | --- | --- | --- | --- |
| alpha Tubulin Monoclonal Antibody (YL1/2) | Rat | Invitrogen (MA1-80017) | IF/SR: 1:1000 | a-tubulin (super-resolution) |
| Anti-CENP-C (Human) pAb (Polyclonal Antibody) | Guinea pig | MBL International (PD030) | IF: 1:1000 | CENP-C |
| Anti-Centromere Protein Antibody | Human | Antibodies Incorporated (15-234) | IF: 1:300 | CREST |
| CENP-A monoclonal antibody (3-19) | Mouse | Enzo Life Sciences (ADI-KAM-CC006-E) | IF: 1:1000 | CENP-A |
| Mouse IgG (H+L) Alexa 488 | Donkey | Jackson Immuno Research (715-545-150) | IF: 1:1000 | NA |
| Mouse IgG (H+L) Alexa 555 | Goat | Thermo fisher Scientific (A-21424) | IF: 1:1000 | NA |
| Mouse IgG (H+L) Alexa 647 | Goat | Thermo fisher Scientific (A-21235) | IF: 1:1000 | NA |
| Mouse IgG (H+L) CF770 | Goat | Biotium (20077) | WB: 1:10000 | NA |
| PP2A A Subunit (81G5) Rabbit Monoclonal Antibody #2041 | Rabbit | Cell Signalling Technology (2041T) | WB: 1:1000 | PPP2R1A |
| Rabbit anti-KIF18A Antibody | Rabbit | Bethyl Laboratories (A301-080A-T) | IF: 1:1000 | KIF18A |
| Rabbit IgG (H+L) Alexa 488 | Goat | Thermo fisher Scientific (A-11034) | IF: 1:1000 | NA |
| Rabbit IgG (H+L) Alexa 555 | Goat | Thermo fisher Scientific (A-21429) | IF: 1:1000 | NA |
| Rabbit IgG (H+L) Alexa 647 | Goat | Thermo fisher Scientific (A-21245) | IF: 1:1000 | NA |
| Rabbit IgG (H+L) CF680 | Goat | Thermo fisher Scientific (A-21434) | WB: 1:10000 | NA |
| α-Tubulin | Mouse | Sigma (T6199) | IF/WB: 1:1000 | a-tubulin |

**Supplemental Table 6:**
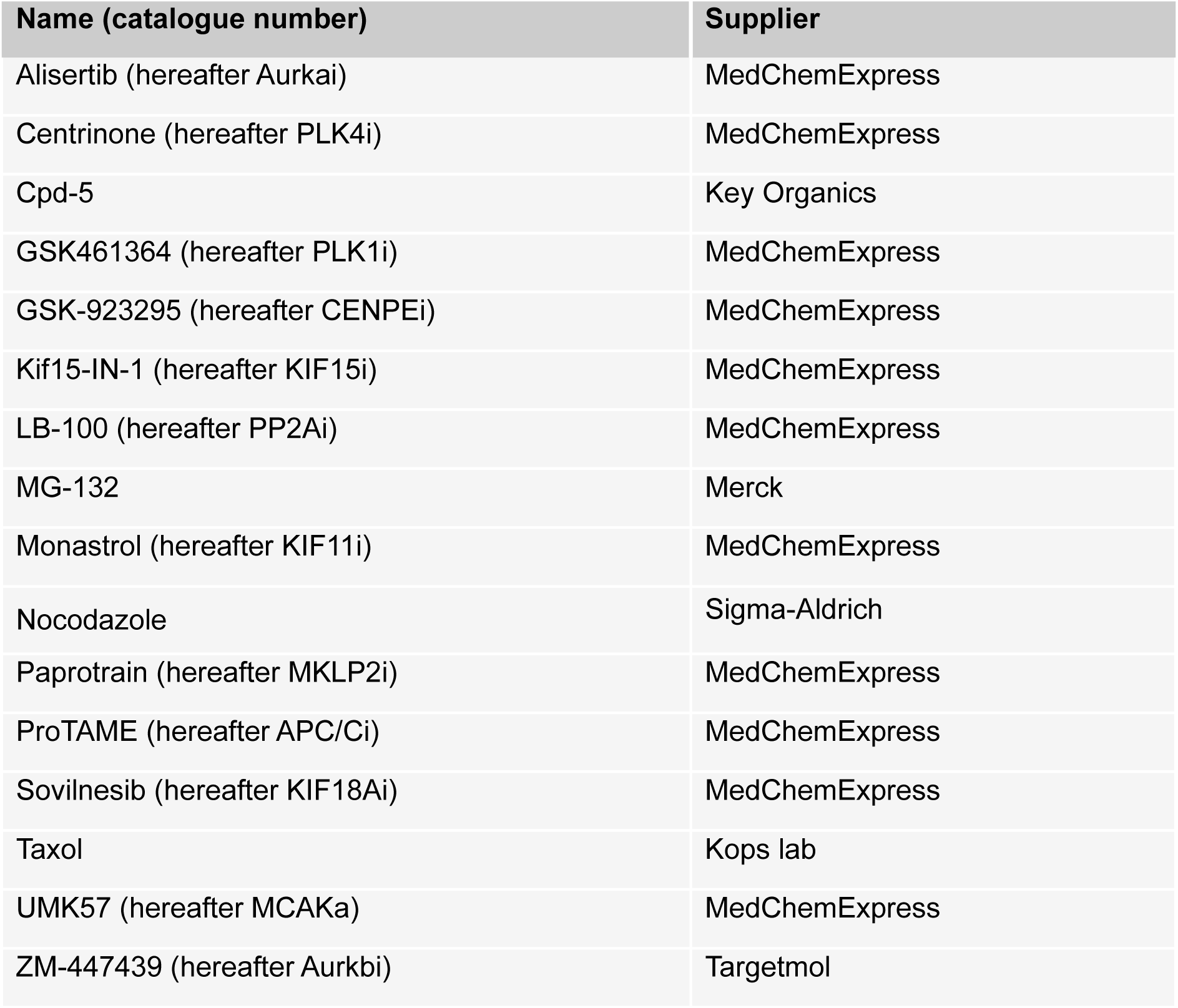
Compounds.

| **Name (catalogue number)** | **Supplier** |
| --- | --- |
| Alisertib (hereafter Aurkai) | MedChemExpress |
| Centrinone (hereafter PLK4i) | MedChemExpress |
| Cpd-5 | Key Organics |
| GSK461364 (hereafter PLK1i) | MedChemExpress |
| GSK-923295 (hereafter CENPEi) | MedChemExpress |
| Kif15-IN-1 (hereafter KIF15i) | MedChemExpress |
| LB-100 (hereafter PP2Ai) | MedChemExpress |
| MG-132 | Merck |
| Monastrol (hereafter KIF11i) | MedChemExpress |
| Nocodazole | Sigma-Aldrich |
| Paprotrain (hereafter MKLP2i) | MedChemExpress |
| ProTAME (hereafter APC/Ci) | MedChemExpress |
| Sovilnesib (hereafter KIF18Ai) | MedChemExpress |
| Taxol | Kops lab |
| UMK57 (hereafter MCAKa) | MedChemExpress |
| ZM-447439 (hereafter Aurkbi) | Targetmol |

