## Supplemental Figures for "Mitotic adaptations shape acquired resistance and vulnerabilities to KIF18A inhibition in cancer"

Figure S1

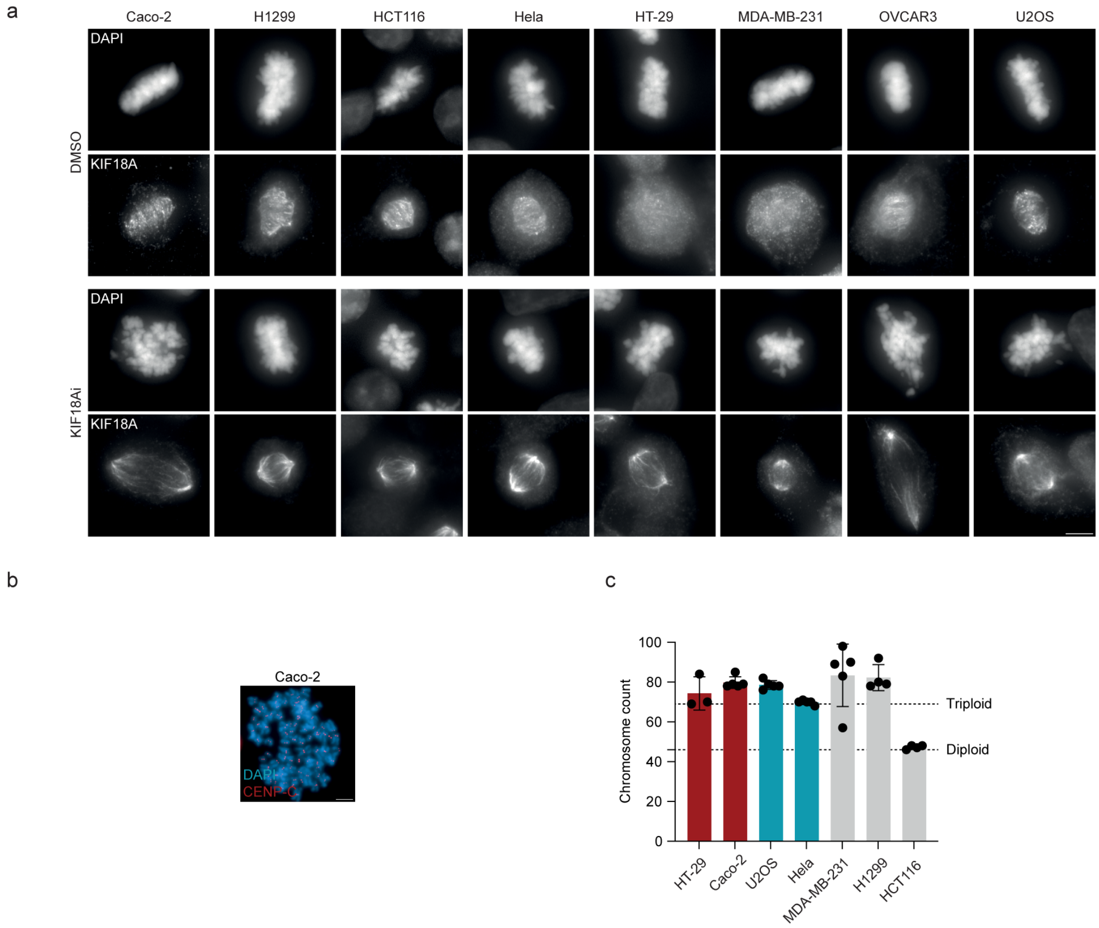


Figure S1. Characterization of cancer cell line panel.

(a) Localization of KIF18A after KIF18Ai in cancer cell line panel using immunofluorescence.

(b) Example image, and

(c) quantification of chromosome counts after performing metaphase spreads. Scale bar = 5 μm.

Figure S2

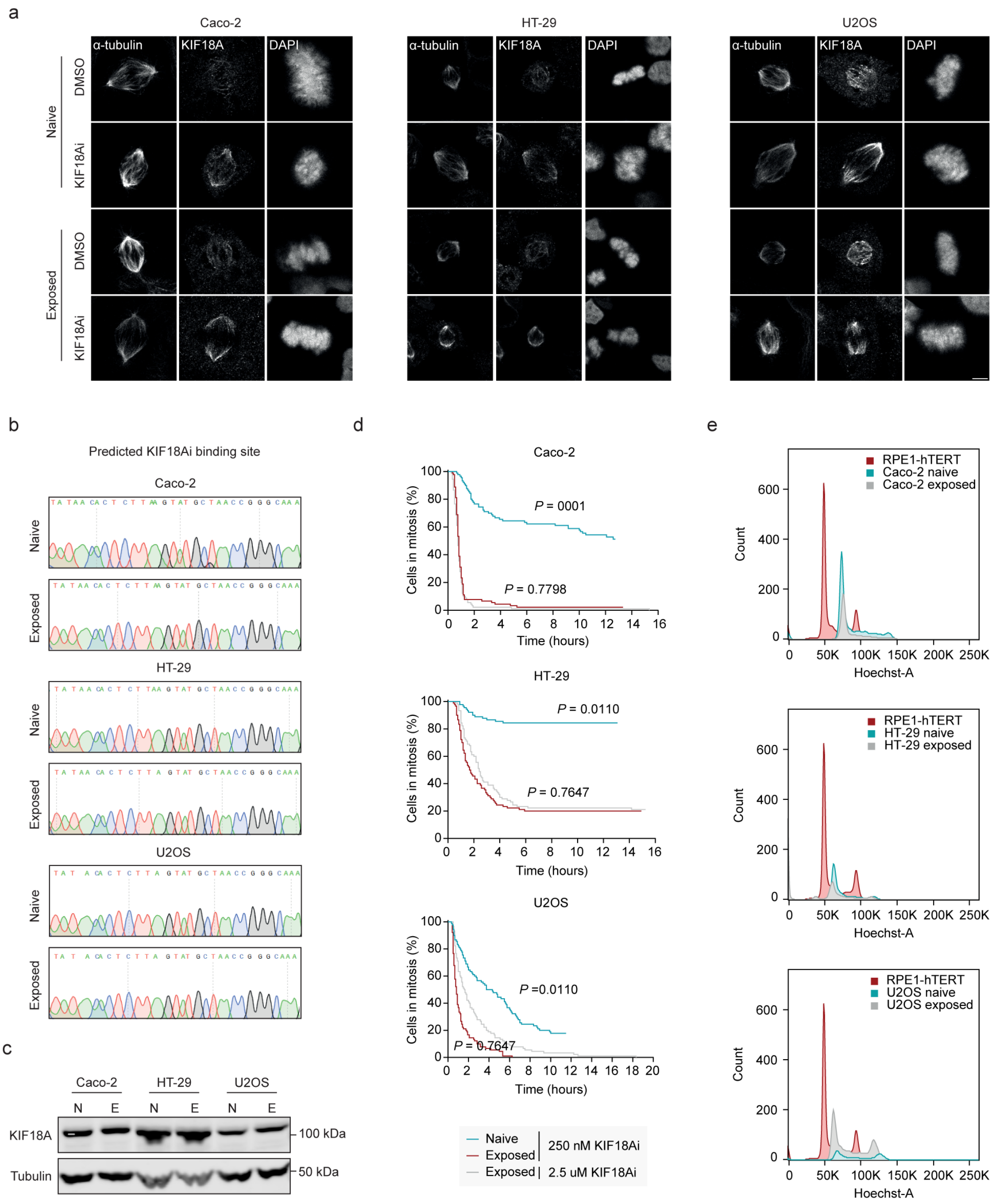
a
Caco-2
HT-29
U2OS
α-tubulin
KIF18A
DAPI
Naive
Exposed
DMSO
KIF18Ai
b
Predicted KIF18Ai binding site
Caco-2
HT-29
U2OS
Naive
Exposed
c
Caco-2
HT-29
U2OS
N
E
KIF18A
Tubulin
100 kDa
50 kDa
d
Caco-2
P = 0001
P = 0.7798
HT-29
P = 0.0110
P = 0.7647
U2OS
P =0.0110
P = 0.7647
Cells in mitosis (%)
Time (hours)
Naive
Exposed
Exposed
250 nM KIF18Ai
2.5 uM KIF18Ai
e
RPE1-hTERT
Caco-2 naive
Caco-2 exposed
HT-29 naive
HT-29 exposed
U2OS naive
U2OS exposed
Count
Hoechst-A

Figure S2. KIF18A is still inhibited and ploidy is unaltered in exposed cancer cells.

(a) Representative immunofluorescence images of metaphases in naïve and exposed cancer cell lines treated with DMSO or KIF18Ai. Scale bar = 5 µm.

(b) Example Sanger sequencing results of the part of the gene encoding for the predicted KIF18Ai interaction site. The full KIF18A gene was sequenced.

(c) Representative western blot of KIF18A protein levels in naïve and exposed cells. Tubulin was used as a loading control. Experiment was performed in triplicate.

(d) Time in mitosis for naïve and exposed cells after treatment with normal (250 nM) or high (2.5 µM) concentrations of KiF18Ai. Three independent experiments were performed (one-way ANOVA with Dunnett's multiple comparisons test using 2.5 µM KIF18Ai as the control; *n* = 90).

(e) Flow cytometry results of the DNA content of indicated cell lines. RPE1-hTERT $TP53^{-/-}$ cells were used as a diploid control.

Figure S3

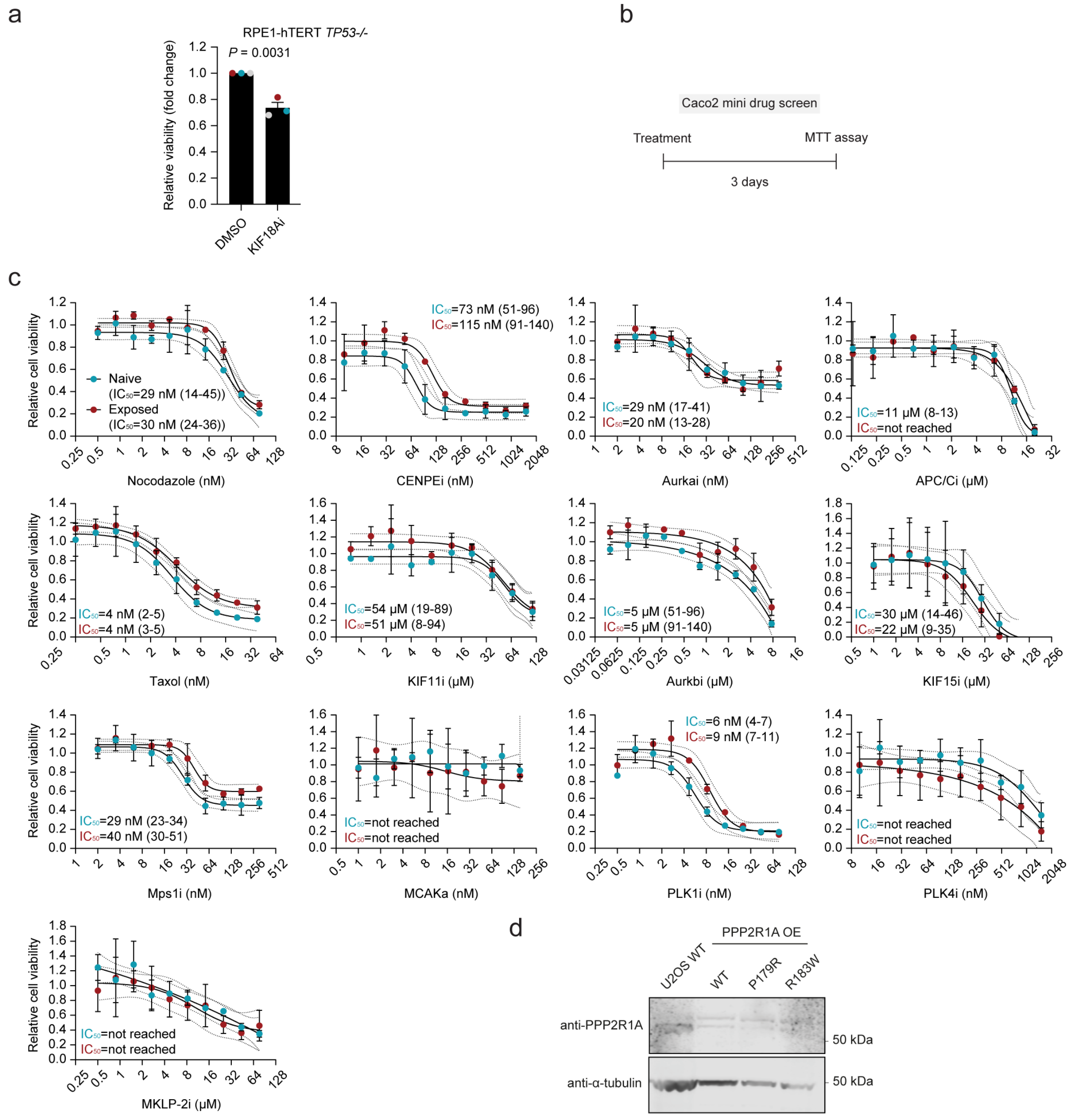


Figure S3. Resistant cancer cells do not become sensitive to other mitotic drugs.
(a) CellTiter-Glo assay was performed in triplicate (mean ± s.d., two-way unpaired t-test).
(b) Schematic illustration of the drug screen.
(c) Quantifications of the cell viability using an MTT assay after a 3 day treatment with indicated drugs. Two independent experiments were performed in technical duplicate (mean ± s.d., dose-response curve based on four-parameter logistic model with 95% confidence interval).
(d) Western blot of U2OS cells overexpressing flag-tagged PPP2R1A after lentiviral infection. Staining was performed using PPP2R1A antibody. Tubulin was used as a loading control.
